# Self-Assembly Coupled to Liquid-Liquid Phase Separation

**DOI:** 10.1101/2022.10.13.512015

**Authors:** Michael F. Hagan, Farzaneh Mohajerani

**Affiliations:** Martin A. Fisher School of Physics, Brandeis University, Waltham, MA, 02453

## Abstract

Liquid condensate droplets with distinct compositions of proteins and nucleic acids are widespread in biological cells. While it is known that such droplets can regulate irreversible protein aggregation, their effect on reversible self-assembly remains largely unexplored. In this article, we use kinetic theory and solution thermodynamics to investigate the effect of liquid-liquid phase separation on the reversible self-assembly of structures with well-defined sizes and architectures. We find that when assembling subunits preferentially partition into liquid domains, robustness against kinetic traps and maximum achievable assembly rates can be significantly increased. In particular, the range of solution conditions over which productive assembly and the corresponding assembly rates can increase by orders of magnitude. We analyze the rate equation predictions using simple scaling estimates to identify effect of liquid-liquid phase separation as a function of relevant control parameters. These results may elucidate self-assembly processes that underlie normal cellular functions or pathogenesis, and suggest strategies for designing efficient bottom-up assembly for nanomaterials applications.

## I. INTRODUCTION

The self-assembly of basic subunits into larger structures with well-defined architectures underlies essential functions in biological organisms, where examples of assembled structures include multi-protein filaments such as microtubules or actin [1, 2], scaffolds for vesicular budding [3–9] the outer shells or ‘capsids’ of viruses [10–16], and bacterial microcompartments [17–22] or other proteinacious organelles [23–27]. However, achieving efficient and high fidelity assembly into target architectures requires precisely tuned subunit interaction strengths and concentrations due to competing thermodynamic and kinetic effects (e.g. [28–47]). The need for such precision could severely constrain the use of assembly for biological function or human engineered applications. Biological organisms employ multiple modes of biochemical and physical regulation to overcome this limitation. In this article, we investigate one such mode — how spatial heterogeneity due to formation of biomolecular condensates can dramatically enhance the speed and robustness of self-assembly.

While membranous organelles play a prominent role in compartmentalizing eukaryotic cells, it is now clear that condensates act as ‘membrane-less’ organelles to spatially organize cellular interiors within all kingdoms of life (e.g. [48–69]. These domains are implicated in diverse cellular functions, including transcriptional regulation [53, 70– 73], formation of neuronal synapses [74–76],enrichment of specific proteins and nucleic acids [77–82], cellular stress response [83–86], and cell division [79, 87]. In addition to the roles of condensates in normal cellular function, pathogenic viruses generate or exploit liquid domains during various stages of their life cycles [88–93]. Most relevant to this article, many viruses undergo assembly and/or genome packaging within phase-separated domains known as virus factories, replication sites, Negri bodies, inclusion bodies, or viroplasms [88–108]. *In vitro* studies show that viral nucleocapsid protein and RNA undergo LLPS (e.g. [99, 106, 109–111], and that LLPS accelerates assembly of nucleocapsid-like particles [99]. It is hypothesized that viruses exploit LLPS to avoid host immune responses and coordinate events such as RNA replication, capsid protein translation, assembly, and genome packaging. However, the mechanisms underlying these events are poorly understood.

Multiple lines of evidence suggest that condensate formation is driven by favorable interactions among their constituents combined with unfavorable interactions with the bulk exterior cytoplasm or nucleoplasm. Although condensation may be driven, destabilized, or regulated by diverse nonequilibrium effects (e.g. [54, 70–73, 112–115], equilibrium thermodynamics provides a starting point to model their stability, and their formation is frequently described as LLPS [48, 49, 52–60, 69, 74, 85, 112–122]. Henceforth, we will use the term LLPS, keeping in mind that nonequilibrium effects may also be present. Consistent with equilibrium phase coexistence, the composition of the domain interior can significantly differ from that of the cytoplasm. Thus, LLPS can provide significant spatiotemporal control over reaction processes by concentrating and colocalizing specific sets of subunit species that preferentially partition into the domain.

These capabilities potentially enable LLPS to strongly regulate self-assembly. Yet, despite recent intense investigations into LLPS, its coupling to assembly has yet to be fully explored. Previous simulations showed that the condensed enzyme complex that forms the interior cargo of bacterial microcompartments can promote nucleation and control the size of the exterior protein shell [43, 45, 123–125]. Most closely related to our work, Refs. [126–129] recently showed that the presence of a domain can significantly accelerate irreversible protein aggregation into linear fibrils.

Here, we investigate the effects of LLPS on *reversible* self-limited assembly into target structures with finite sizes and well-defined architectures. Self-limited assembly from bulk solution is constrained by competing thermodynamic and kinetic effects — subunit interactions must be sufficiently strong and geometrically precise to stabilize the target structure, but overly high interaction strengths or subunit concentrations lead to kinetic traps [28–47]). Avoiding such kinetic traps imposes a ‘speed limit’ on assembly from bulk solution [41, 42, 130].

Using a master equation description of assembly, we show that these thermodynamic and kinetic constraints can be simultaneously satisfied by spatial heterogeneity due to phase-separated domains. We find that LLPS can significantly accelerate assembly nucleation, consistent with previous studies of irreversible assembly [126–129], but also induces kinetic traps that *slow* assembly in certain parameters regimes. Crucially though, by enhancing nucleation only within spatially localized regions, LLPS significantly expands the range of subunit concentrations and interaction strengths over which such kinetic traps are avoided, thus promoting assembly robustness. This effect can increase by orders of magnitude the maximum rate of *productive* assembly into the ordered target structure. The extent of assembly acceleration and robustness enhancement are nonmonotonic functions of the key control parameters: the partition coefficient of subunits between the domain and bulk cytoplasm and the domain volume. We present simple scaling estimates that qualitatively capture the effect of LLPS on assembly, and reveal the underlying mechanisms that enable regulation. For example, the bulk solution acts as a “buffer” that steadily supplies free subunits to the domain to enable rapid assembly without kinetic traps. Although we particularly focus on self-limited assembly processes that lead to finite-sized structures, our models are general and many results also apply to unlimited assembly or crystallization.

## II. RESULTS AND DISCUSSION

### A. Model

We have developed a minimal model to describe assembly in the presence of one or more liquid droplets coexisting with a background solution of different composition. We are motivated by processes such as virus assembly, in which viral proteins, nucleic acids and other viral components, and possibly some host proteins phase separate to form liquid domains within the cellular cytoplasm. For this initial study, we consider only one assembling species in the limit that the assembly subunits comprise a small fraction of the domain mass, and thus the size and composition of the domain can be treated as independent of the subunit concentration.

We consider a system of subunits that form self-limited assemblies with optimal size *N*. The subunits are immersed in a binary solvent (which could be proteins, polymers, nucleic acids, etc.) which is in a state of phase coexistence, with stoichiometry such that there are one or more small domains rich in one solvent species coexisting with a much larger background rich in the other species. We denote the volumes of the domain and background as *V*_dom_ and *V*_bg_, which are related to the total system volume by *V*_tot_ = *V*_dom_ + *V*_bg_. We will present results in terms of the domain size ratio, *V*_r_ ≡ *V*_dom_*/V*_bg_. In most biological systems or in vitro experiments, the domain volume will be small compared to the background, *V*_r_ ≪ 1. For this work we assume a fixed total subunit concentration *ρ*_T_. For simplicity we will typically consider a single domain, but we also discuss the case of multiple domains, which might arise due to microphase separation or arrested phase separation.

The subunits preferentially partition into the domain with a partition coefficient *K*_dom_, which at equilibrium satisfies

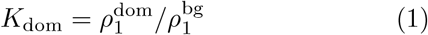

with 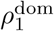 and 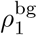 the subunit concentrations in the domain and background.

The partition coefficient is related to the change in solvation free energy *g*_dom_ for a subunit that transitions from the background to the domain as 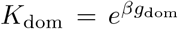 with *β* = 1*/k*_B_*T* with *k*_B_*T* the thermal energy. Applying standard dilute solution thermodynamics will result in a subunit solvation free energy difference with the form (see Weber et al. [114]) *g*_dom_ ∝ *n*_s_(Δ*χ*)(Δ*ϕ*) where Δ*ϕ* is the difference in solvent composition between the background and domain, Δ*χ* is the difference in interaction strength (parameterized by the Flory *χ* parameter) for a interaction site on the subunit between the background and domain, and *n*_s_ is the number of subunit interaction sites. The key point is that even for relatively weak interactions, a multivalent subunit with *n*_s_ ≳ 10 could have a partition coefficient as large as *K*_dom_ ∼ 10^4^ − 10^5^, although *K*_dom_ ∼ 100 may be a typical value [54].

### B. Effects of LLPS on self-assembly equilibrium

We begin by calculating how the equilibrium yield of self-assembled structures depends on subunit concentrations and interaction strengths, as well as the two key control parameters for subunit partitioning into the domain: the partition coefficient *K*_dom_ and the domain size ratio *V*_r_.

At equilibrium the subunit concentrations in the domain and background are related to each other by *K*_dom_, and to the total subunit concentration *ρ*_T_ by mass conservation, giving

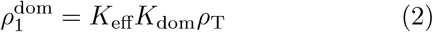

with

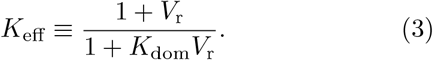

#### Assembly yield and spatial control over assembly

We now calculate the effect of the domain on assembly yields, using the well-justified approximation that intermediates have very low concentrations at equilibrium for self-limited assembly [41, 131]. Thus we consider a two-state system, with finite concentrations of only free subunits and complete assemblies with *N*. Mass conservation then gives

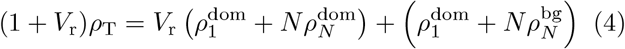

where 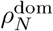 and 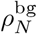 are the concentrations of assemblies in the domain and background. At equilibrium these are related to the free subunit concentration by the law of mass action [41, 132]

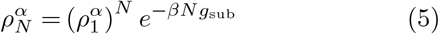

with *α* = dom, bg and *g*_sub_ as the per-subunit interaction energy within a complete assembly (which we assume is the same in the domain and background). Eqs. (4), (5) can be easily solved numerically. However, we can simplify the analysis by assuming the limit of assembly only occurring within the domain, so 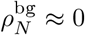. Then, we write the fraction of subunits in assemblies as

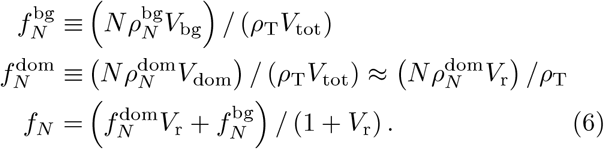

We then substitute 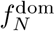 into Eq. (4), and following the analysis for assembly of capsids in [41], we obtain

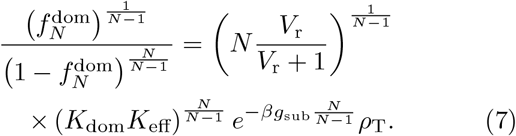

In the limit of large optimal assembly size *N* ≫ 1, Eq. (7) satisfies the following asymptotic limits

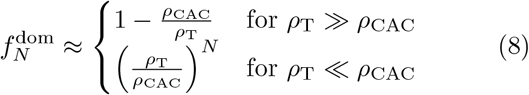

with *ρ*_CAC_ the *critical assembly concentration* (CAC) given by

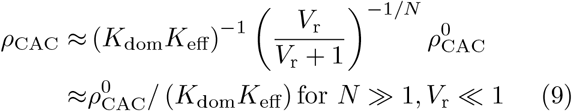

with

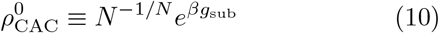

as the CAC in a system without coupling to LLPS (i.e. *V*_r_ = 0 or *K*_dom_ = 1). In all subsequent expressions, we will write the limit of no LLPS or *K*_dom_ → 0 with a superscript ‘0’. The last expression in Eq. (9) assumes *V*_r_ ≪ 1 and shows that LLPS reduces the CAC by a factor *K*_dom_*K*_eff_. Then using Eq. (2) we arrive at the simple result that significant assembly occurs when the *total* subunit concentration *ρ*_T_ exceeds the *local* CAC within the domain.

To obtain further insight, we note that *K*_dom_*K*_eff_ ≈ (*V*_r_ + 1*/K*_dom_)^*−*1^, giving the asymptotic limits

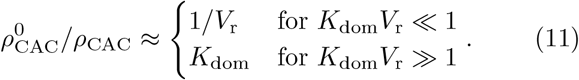

and that maximal enhancement of equilibrium assembly is achieved when *K*_dom_ ≳ *V*_r_.

We can draw two important conclusions from Eq. (9). First, the presence of a domain allows assembly under conditions where there is no bulk assembly (Fig. 2). Second, there is a range of total subunit concentrations ∝ *K*_dom_*K*_eff_ over which assembly occurs only in the domain, thus allowing for spatial control over assembly. As a measure of the extent to which LLPS can spatially control assembly, we define *selectivity* as 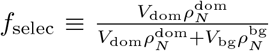

**FIG. 1.**
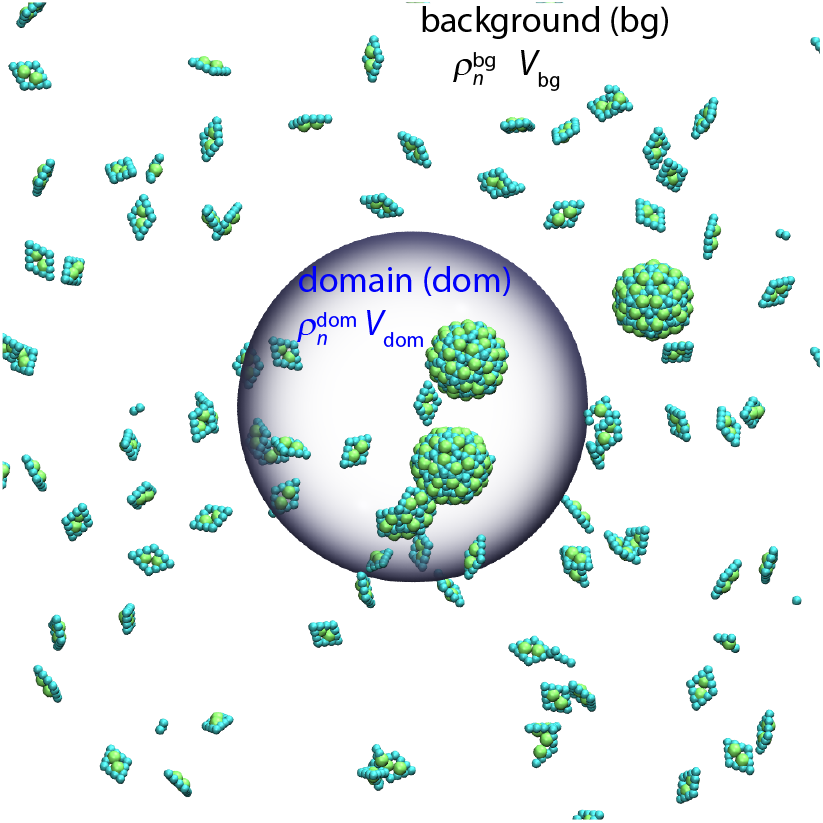
Schematic of the model. Subunits exchange between bulk and the phase-separated domain (gray sphere), with equilibrium concentrations related by 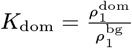. Assembly can occur anywhere in the system, but occurs preferentially in the domain when *K*_dom_ *>* 1 due to the enhanced local subunit concentration. The volumes of the bulk *V*_bg_ and domain *V*_dom_ are related by *V*_r_ = *V*_dom_*/*(*V*_dom_ + *V*_bg_).

**FIG. 2.**
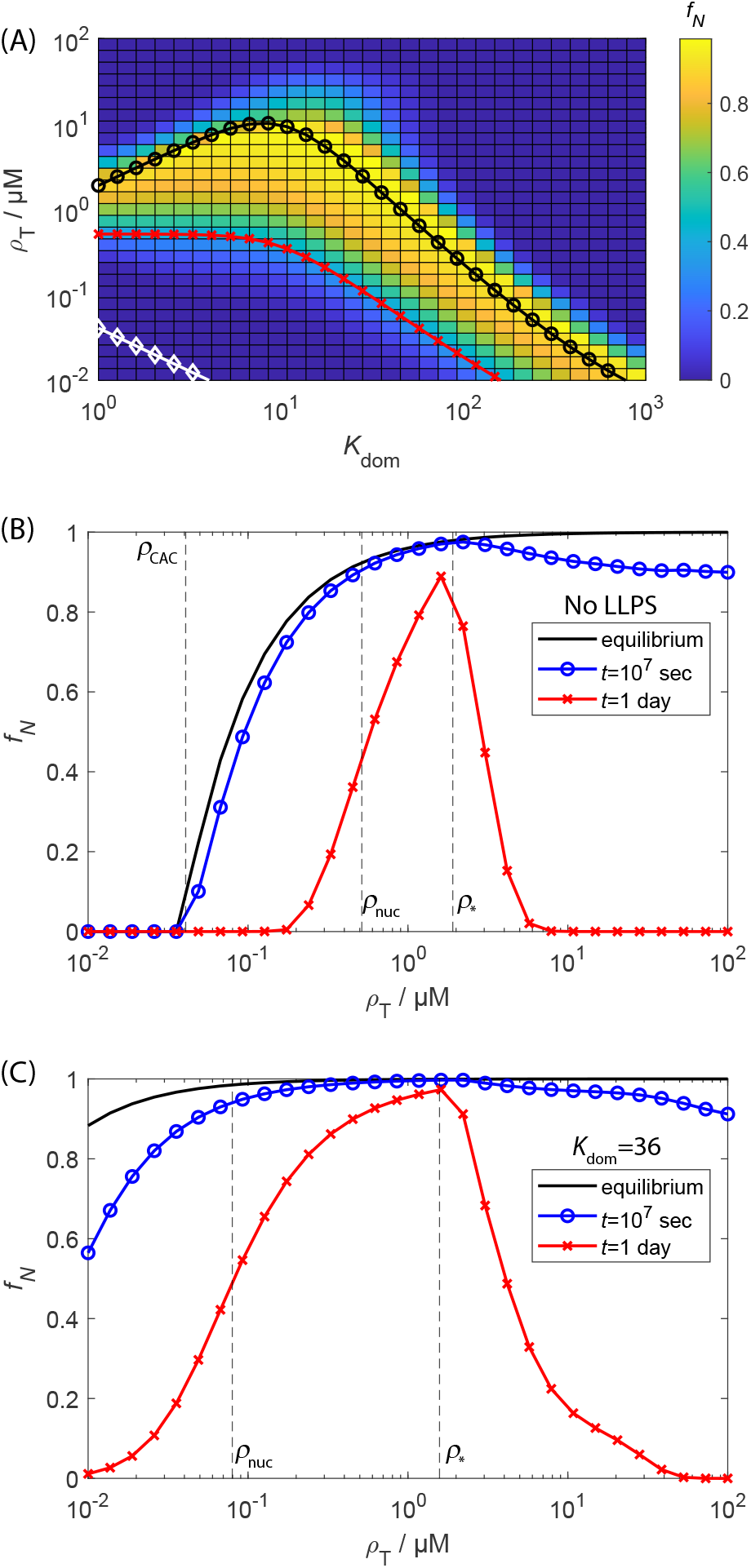
Effect of LLPS on the equilibrium and finite-time yields of self-assembly. **(A)** The heat map shows the mass fraction of subunits in capsids *f*_*N*_ as a function of the domain partition coefficient *K*_dom_ and total subunit concentration *ρ*_T_ computed from the rate equations with the nucleation-andgrowth (NG) model (Eq. (13)) at a finite time of 1 day. The lines show: the equilibrium critical assembly concentration (*ρ*_CAC_, Eq. (9), white ‘⋄’ symbols), the predicted threshold parameter values below which the median assembly timescale *τ*_1*/*2_ exceeds 1 day (*ρ*_nuc_, Eq. (28), red ‘x’ symbols), and the predicted locus of points corresponding to the minimum assembly timescale, beyond which monomer starvation begins to set in (*ρ*_*_, Eq. (31), black ‘◦’ symbols). **(B)** The mass fraction of complete capsids *f*_*N*_ as a function of total subunit concentration for no LLPS (*K*_dom_ = 1). The line shows the equilibrium result (Eq. (7)) and the symbols show results from numerically integrating the rate equations to 1 day (∼ 9 × 10^4^ sec, blue ‘⋄’ symbols) and *t* = 10^7^ seconds (red x symbols). The dashed lines show *ρ*_CAC_, *ρ*_nuc_, and *ρ*_*_. **(C)** Same as (B), but in the presence of LLPS, with *K*_dom_ = 36. Other parameters in (A-C) are critical nucleus size *n*_nuc_ = 3, optimal size *N* = 120, subunit binding affinities *g*_nuc_ = −4*k*_B_*T, g*_elong_ = −17*k*_B_*T*, and domain volume ratio *V*_r_ = 10^*−*3^.

The equilibrium selectivity is then given by

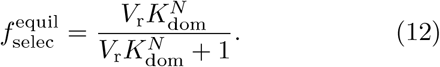

We thus see that even a very small partition coefficient leads to strong equilibrium selectivity due to the highvalence nature of an assembled capsid. In particular, an assembled capsid has ∼ *N* interactions with domain components, but only has three translational degrees of freedom suppressed by partitioning into the domain volume. However, if assembled capsids and large intermediates are not able to rapidly exchange between the domain and background [129], the selectivity at finite-times may be under kinetic control.

### C. Effect of LLPS on self-assembly kinetics

#### 1. Master equation models for capsid assembly

To simulate the assembly kinetics, we adapt the rate equation description originally developed by Zlotnick and coworkers [28, 29, 133] and used by others [130, 134] to describe the self-assembly of 2D polymers (capsids) in bulk solution. Denoting the concentration of an intermediate with *n* subunits in either phase as 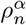 with *α* = dom,bg, the time evolution of intermediate concentrations is given by:

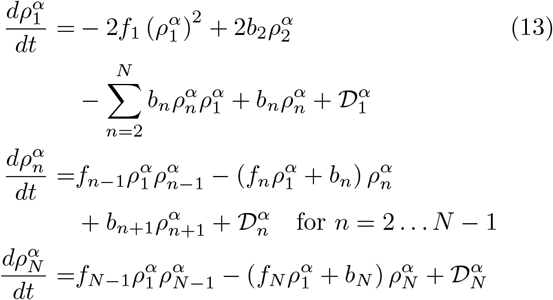

with the diffusive exchange between the phases given by (see Appendix A and Refs. [126, 129])

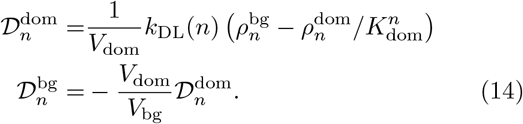

and *f*_*n*_ and *b*_*n*_ as the association and dissociation rate constants for intermediates of size *n*. Since we have assumed low concentrations of subunits in the domain, we assume that the diffusion coefficient is independent of subunit concentration. To focus on effects of competing reactions on assembly, we also neglect the possible dependence of diffusion coefficients on intermediate size or *g*_dom_, considered in Refs. [128, 129] respectively. The model can be readily extended to account for these effects.

We set the initial condition as 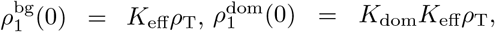 and 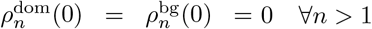

##### Nucleation and growth model (NG)

To complete the Master equation description we must specify the set of states, which in this case corresponds to the set of allowed assembly intermediates and products, and their associated transition rates. We start with a simple generic model for a nucleated self-assembly process denoted as the ‘nucleation and growth (NG) model’ [130]. This can describe linear assembly with nucleation (e.g. assembly of a helical viral capsid or the equilibrium assembly of an actin filament), or polyhedral shell assembly with an initial nucleation step, followed by assembly along a single growth front until the shell closes on itself [29, 130, 134]. We consider a system of capsid protein subunits with total concentration *ρ*_T_ that start assembling at the time *t* = 0. We assume that the rate constants are the same in the domain and background, so we simplify the presentation by omitting the specification of phase in this subsection. Our reaction is given by:

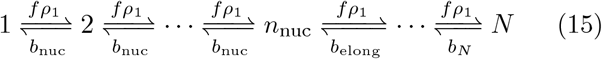

where *b*_*i*_ is the dissociation rate constant (with *i* = {nuc, elong, *N*}), which is related to the forward rate constant by detailed balance as *v*_0_*b*_*i*_ = *f* exp (*βg*_*i*_), with *g*_*i*_ the change in interaction free energy upon subunit association to a partial capsid and *v*_0_ the standard state volume. The nucleation and elongation phases are distinguished by the fact that association in the nucleation phase has an unfavorable free energy change, *g*_nuc_ −*k*_B_*T* ln (*ρ*_1_*v*_0_) > 0, while association in the elongation phase is favorable, *g*_elong_ − *k*_B_*T* ln (*ρ*_1_*v*_0_) < 0. For the moment, we assume that there is a single critical nucleus size *n*_nuc_.

For most results in this article, we will set *g*_nuc_ = −4*k*_B_*T* and *g*_elong_ = −17*k*_B_*T* and *g*_N_ = 2*g*_elong_. The small value of *g*_nuc_ relative to *g*_elong_ accounts for the fact that the first few subunits to associate make fewer and/or less favorable contacts than subunits in larger intermediates, giving rise to a nucleation barrier, while the large value of *g*_N_ accounts for the fact that in many assembly geometries the last subunit makes the largest number of contacts upon associating. For concreteness, we have chosen the specific values of *g*_nuc_ and *g*_elong_ = −17*k*_B_*T* to be roughly consistent with binding affinity values estimated for virus capsid assembly [28, 30, 31, 135], but the results do not qualitatively change for other affinity values within a given assembly regime.

##### Classical nucleation theory model (CNT)

To test whether our conclusions depend qualitatively on the model geometry, we also consider transition rates based on the ‘classical nucleation theory (CNT)’ model for icosahedral capsids suggested by Zandi et al. [136]. In this model, assembly intermediates are represented as partial spheres that are missing a spherical cap. Subunits along the perimeter of the missing cap have fewer interactions than those in the shell interior, leading to a line tension *σ*, and the binding free energy is

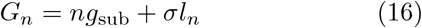

with the perimeter of the missing spherical cap given by

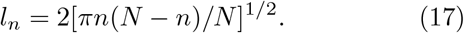

with *g*_sub_ the binding free energy per subunit in a complete capsid. Following previous work [130, 136], we set the line tension to *σ* = − *g*_sub_/2, so that a subunit adding to the perimeter of the capsid satisfies half of its contacts on average. We assume that the forward rate constant is proportional to the number of subunits on the perimeter, *f*_*n*_ = *f*_0_*l*_*n*_, with *f*_0_ the association rate constant for a single binding site.

The key difference between the CNT and NG models is the dependence of the critical nucleus size on solution conditions. For the NG model the critical nucleus size *n*_nuc_ is constant, provided exp(*g*_nuc_*/k*_B_*T*) < *ρ*_1_*v*_0_ < exp(*g*_elong_*/k*_B_*T*). For the CNT model, the critical nucleus size varies with subunit concentration and interaction strengths, and is given by the maximum in *k*_B_*T* log(*ρ*_1_*v*_0_)*n* + *G*_*n*_, or [136]

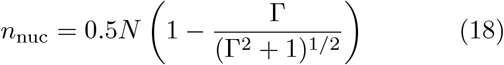

with Γ = [*g*_sub_ ln(*ρ*_1_*v*_0_)]*/σ*. Thus, the critical nucleus size continually changes over time during an assembly process for the CNT model as subunits are depleted, whereas it is constant until the very late stages of a NG assembly process.

#### 2. Master equation results

Figs. 2 and 3 show the effect of LLPS on assembly kinetics, as measured by the fraction of subunits in complete capsids (*f*_*N*_), obtained by numerically integrating the Master equation (Eq. 13) with the NG model (Eq. 15). Fig. 3A shows *f*_*N*_ as a function of time for several initial subunit concentrations *ρ*_T_ in the absence of LLPS. There is an initial lag phase during which intermediate populations build up to a quasi-steady-state, followed by rapid appearance of complete capsids, and then eventually saturation as free subunits are depleted. The duration of the lag phase decreases as 1*/ρ*_T_, and the rate of capsid production is nonmonotonic with respect to *ρ*_T_ — yields of complete capsids are suppressed for *ρ*_T_ = 4 µM by a kinetic trap arising from depletion of free subunits before capsids finish assembling. These results are discussed further in section II D 1 and Refs. [41, 130]. Figs. 3B and 2A show the how the assembly kinetics is changed by LLPS. With the lowest concentration shown in Fig. 3A (*ρ*_T_ = 0.2 µM), *f*_*N*_ is shown as a function of time for increasing values of the partition coefficient *K*_dom_. We see that the yields and assembly rates increase dramatically, with the duration of the lag phase decreasing and the maximum rate of capsid production (corresponding to the nucleation rate) increasing with *K*_dom_. To give a more comprehensive picture, Fig. 2A shows *f*_*N*_ as a function of both *K*_dom_ and *ρ*_T_. We see that assembly occurs at lower concentrations as *K*_dom_ increases, and that LLPS increases the range of concentrations over which productive assembly occurs, particularly for *K*_dom_ of 𝒪 (10). Similarly, Figs. 2B and C respectively show *f*_*N*_ as a function of concentration measured at 1 day, 10^7^ seconds, and equilibrium. Both in the presence and absence of LLPS productive assembly at one day occurs over a much narrower range of concentrations than predicted by equilibrium, due to nucleation barriers at small concentrations and kinetic traps at high concentrations. Even at extremely long times the results have not reached full equilibrium due to kinetic traps at high concentrations. However, LLPS significantly broadens the range of concentrations leading to productive assembly at all timescales. The solid lines in Fig. 2A and the dashed lines in Figs. 2B,C indicate the CAC (*ρ*_CAC_, Eq. (9)), and scaling estimates for threshold concentrations below/above which productive assembly is impeded by large nucleation barriers or kinetic traps respectively (see section section II D 1).

**FIG. 3.**
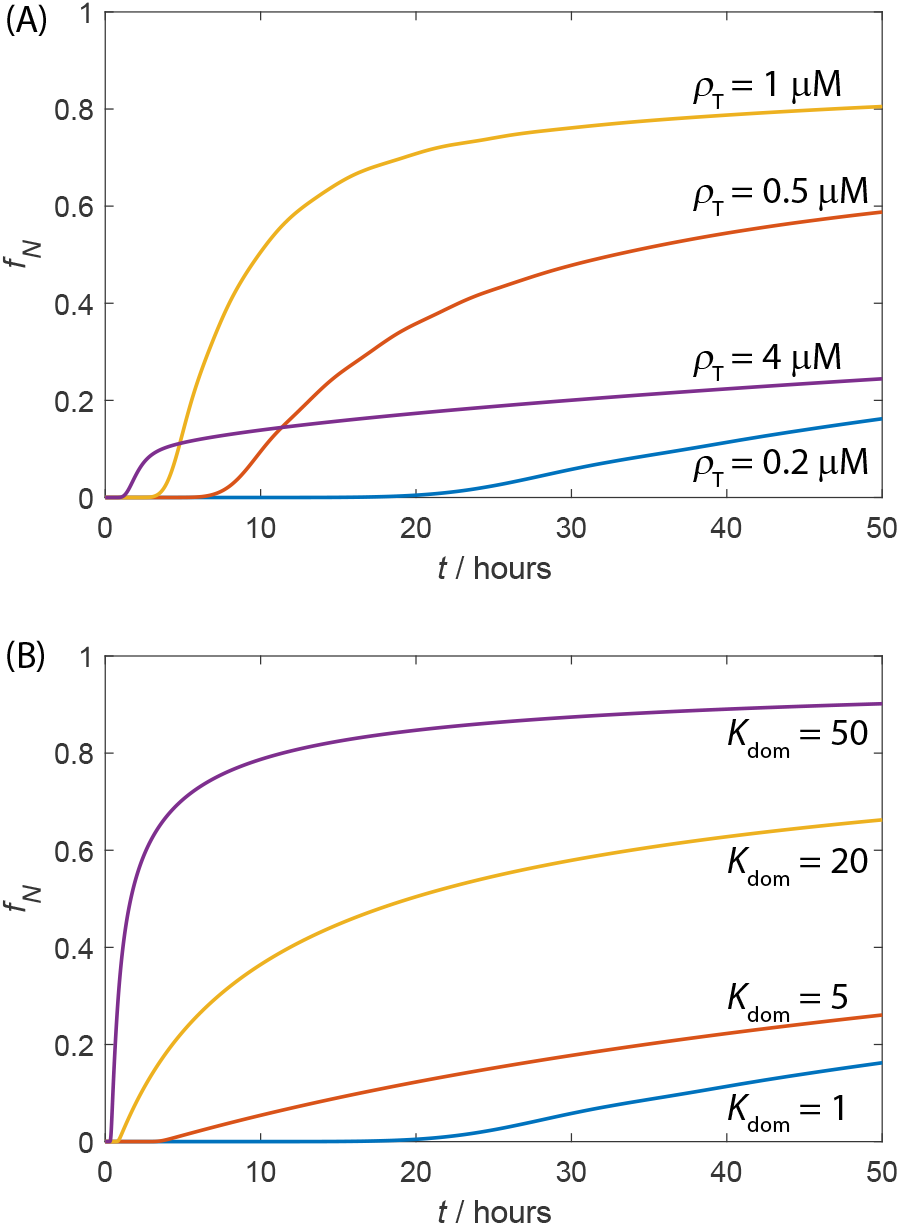
The dependence of assembly kinetics on parameter values for the NG model with and without LLPS. **(A)** The time evolution of the fraction of subunits in complete capsids *f*_*N*_ for indicated values of the total subunit concentration *ρ*_T_ computed from the Master equation, with no LLPS. **(B)** The time evolution of *f*_*N*_ for indicated values of *K*_dom_, for fixed *ρ*_T_ = 0.2*µ*M and domain ratio *V*_r_ = 0.001. Other parameter values for (A) and (B) are optimal assembly size *N* = 120, critical nucleus size *n*_nuc_ = 3, *g*_nuc_ = −4*k*_B_*T, g*_elong_ = −17*k*_B_*T*.

The ability of LLPS to avoid kinetic traps arises because, for *V*_r_ ≪ 1, the background acts like a buffer that steadily supplies free subunit to the domain even when the nucleation rate is large. As a measure of this behavior, Fig. 4 shows the concentration of subunits in the background normalized by the total concentration, 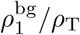 as a function of the maximum capsid formation rate (which occurs shortly after the end of the lag phase, before significant free subunit depletion has occurred). Here we have measured 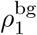 at the time point corresponding to the maximum rate. Results are shown for LLPS assembly for the same parameters as in Fig. 3, with increasing rates corresponding to increasing values of *K*_dom_. For the case without LLPS, we achieve faster rates by increasing the subunit-subunit affinities from (*g*_nuc_, *g*_elong_) = (−4, −17) *k*_B_*T* to (*g*_nuc_, *g*_elong_) = (−6, −25.5) *k*_B_*T*. We have increased affinities rather than total concentration (as we do for other results) to simplify comparison of 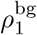 between the two cases. The results stop at *g*_nuc_ = −6 because stronger affinities lead to decreasing rates due to the monomer starvation trap.

**FIG. 4.**
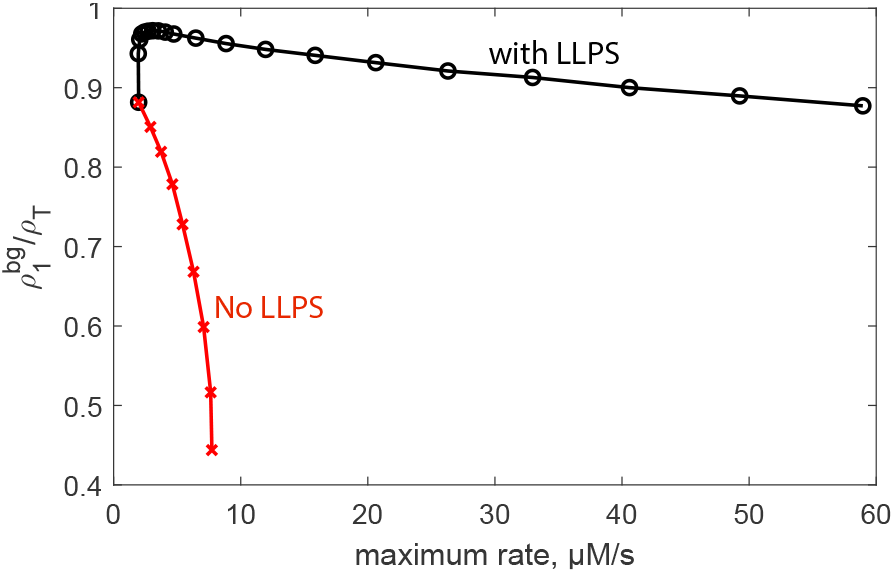
The background acts as a buffer of free subunit for LLPS-dominated assembly. The plot shows the concentration of subunits in the background, 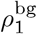, as a function of the maximum capsid formation rate (maximized over time at a given set of parameter values) for assembly with LLPS (black ‘◦’ symbols) and without LLPS (red ‘x’ symbols). For the LLPS case, the parameters correspond to those in Fig. 3 with the increasing rate corresponding to *K*_dom_ ∈ [1, 30]. For the case without LLPS, the parameters are the same except that *K*_dom_ = 1 and the increasing rate is achieved by scaling the subunit-subunit affinities according to *fg*_nuc_ and *fg*_elong_ with *f* ∈ [1, 1.5]. The results stop at *f* = 1.5 because stronger affinities lead to kinetic traps and thus a decreasing maximum rate.

We see that with LLPS the subunit concentration remains near *ρ*_T_ even for extremely high assembly rates, whereas subunits are rapidly depleted without LLPS. For higher subunit affinities |*g*_nuc_| > 6*k*_B_*T* depletion is so rapid that the monomer starvation trap sets in. Note that, in our Master equation description, subunits will eventually be depleted as *f*_*N*_ → 1 even with LLPS, but in reality excluded volume effects (which are neglected in our model) would suppress assembly rates before this point unless complete capsids are expelled from the domain.

Fig. 5 shows the selectivity (*f*_selec_) as a function of *K*_dom_ for several values of target capsid size *N*. (Figs. 5B) closely match the equilibrium result (Eq. (12)), and that even an extremely small partition coefficient *K*_dom_ ≳ 2 is sufficient to drive highly selective assembly in the domain for large *N*.

**FIG. 5.**
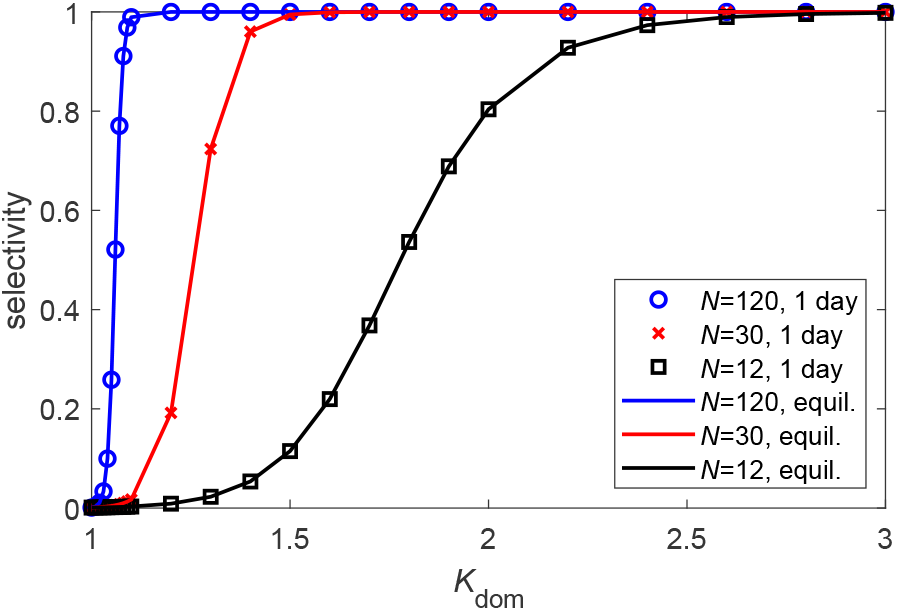
Selectivity as a function of domain partition coefficient and capsid size. The symbols show the selectivity, 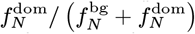 computed from the Master equation as a function of *K*_dom_ for three indicated values of the optimal assembly size *N* at one day. The lines show the equilibrium selectivity (Eq. (12)). Other parameters are domain volume ratio *V*_r_ = 0.001, *g*_nuc_ = 4*k*_B_*T, g*_elong_ = 17*k*_B_*T*, and *ρ*_T_ = 0.2*µ*M.

### D. Scaling Estimates for the Effect Of LLPS on Assembly Timescales

To gain an intuitive understanding of how LLPS can affect assembly, in this section we derive simple scaling estimates for the timescales associated with the nucleation and growth mechanism of Eq. (15). We closely follow Refs. [130, 137], but we extend the analysis to include the effect of a domain. Although we introduce a number of simplifications, in the next section we show that the resulting scaling estimates provide good approximations when these simplifications are relaxed by numerically solving the Master equation models.

#### 1. Timescales without LLPS

Let us begin by summarizing the analysis of Ref [130] for assembly timescales in the absence of LLPS. As above, we consider a system of subunits with total concentration *ρ*_T_ that form assemblies with optimal size *N* subunits, and we break the assembly process into ‘nucleation’ and ‘elongation’ phases. For simplicity we assume that the association rate constant *f* is independent of intermediate size (except where mentioned otherwise), so that the rates of association to each intermediate are *fρ*_1_.

We now write the time required for an individual assemblage to form as *τ* = *τ*_nuc_ +*τ*_elong_ with *τ*_nuc_ and *τ*_elong_ the average times for nucleation and elongation, respectively.

##### Elongation

The elongation timescale can generally be estimated as [130, 131]

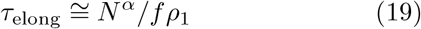

where we have assumed *N* ≫ *n*_nuc_ so that *N* − *n*_nuc_ ≈ *N*. The factor in the numerator indicates that the elongation timescale increases with optimal assembly size (i.e. *α* > 0) since 𝒪 (*N*) independent subunit additions must occur. The value of the exponent *α* will depend on factors such as the dimensionality, the aggregate geometry, and the relative stability of intermediates, but we expect 1/2≤ *α* ≤ 2. For strongly forward-biased assembly during the elongation phase, *α* = 1 for the NG model and *α* = 1/2 for the CNT model (see [130] and Appendix B). Except where specified otherwise, for the scaling estimates in the rest of this article we set *α* = 1, but the results are easily extendable to other exponent values.

##### Nucleation

The mean nucleation time at the beginning of the reaction can be estimated from the statistics of a random walk biased toward disassembly [29, 130], and can be approximately written as

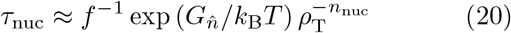

where 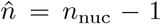 so that 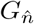 is the interaction free energy of the structure just below the critical nucleus size. The form of Eq. (20) can be understood by noting that the pre-critical nucleus is present with concentration 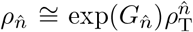, and subunits associate to the precritical nucleus with rate *fρ*_T_. Note that the special case of *n*_nuc_ = 2 corresponds to no nucleation barrier (since two subunits must associate to begin assembly), in which case 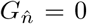 and 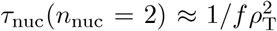. We consider this case in Appendix D.

While Eq. 20 gives the initial nucleation rate, the nucleation rate decreases over time due to subunit depletion, and asymptotically approaches zero as the concentration of completed capsids approaches its equilibrium value. Thus, we estimate the median assembly time *τ*_1*/*2_ (the time at which the reaction is 50% complete) by treating the system as a two-state reaction with *n*_nuc_-th order kinetics, which yields [130]

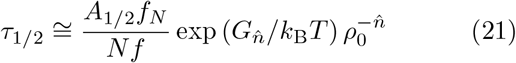

with 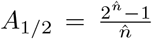, and *f*_*N*_ as the equilibrium fraction of subunits in complete capsids. The factor of *N* ^*−*1^ in Eq. 21 accounts for the fact that *N* subunits are depleted by each assembled capsid.

Analogous to crystallization or phase separation, there is a range of subunit concentrations and interaction strengths for which the unassembled state is metastable; i.e., the system is beyond the CAC so assembly is thermodynamically favorable, but the nucleation timescale exceeds experimentally relevant timescales. The boundary of this regime can be estimated by inverting Eq. (21). Denoting the ‘relevant’ observation timescale as *τ*_obs_, we can estimate the threshold subunit concentration below which nucleation will not be observed as

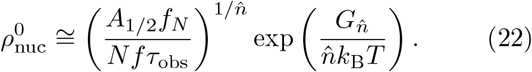

When elongation is fast compared to nucleation, the expressions Eq. B1 and Eq. 21 respectively predict the duration of the lag phase and the median assembly time. However, these relations begin to fail above threshold values of the subunit concentration or subunit-subunit binding affinity, when nucleation and elongation timescales become comparable. Upon further increasing these parameters, nucleation becomes sufficiently fast that a significant fraction of monomers are depleted before elongation of most structures can complete. Subsequent evolution into complete assemblages then requires exchange of subunits between different intermediates (Ostwald ripening), which is an activated process and thus low on assembly timescales. We describe this condition as the *monomer-starvation* kinetic trap. The threshold subunit concentration *ρ*_*_ and interaction energies beyond which the system begins to enter the trap can be estimated by the locus of parameter values at which the median assembly time and elongation time are equal, i.e., *τ*_1*/*2_(*ρ*_*_) = *τ*_elong_(*ρ*_*_):

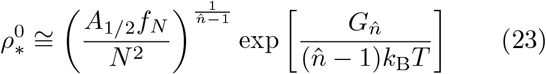

and a corresponding assembly timescale

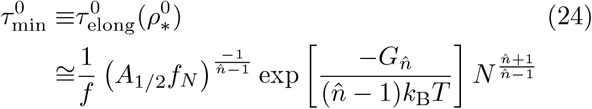

Note that 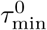 corresponds to approximately the minimal timescale or maximal assembly rate (over all subunit concentrations) since both *τ*_1*/*2_ and *τ*_elong_ monotonically decrease with subunit concentration before the onset of kinetic trapping.

### 2. Assembly timescales with LLPS

We now extend the scaling analysis to account for the presence of a domain. Based on the conclusion of Appendix A that exchange of subunits between the background and domain is typically much faster than assembly rates, we will make a quasi-equilibrium approximation for the relationship between subunit concentrations in the domain and backround: 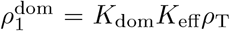 and 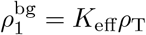

#### Nucleation

As shown previously for irreversible aggregation [114], the domain can dramatically amplify the nucleation rate by locally concentrating subunits. The total initial nucleation rate (in both the domain and background at the beginning of the assembly process) is given by

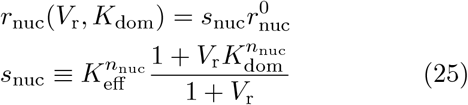

with 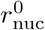 the nucleation rate in the absence of a domain, and *s*_nuc_ the acceleration factor for the initial nucleation rate. Eq. (25) shows that for 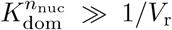 nucleation will occur exclusively in the domain, and the nucleation acceleration factor simplifies to 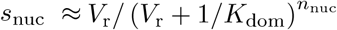.

To estimate the parameters that maximize the initial nucleation rate, we optimize Eq. (25) with respect to to obtain 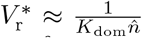 and thus a maximum nucleation acceleration of:

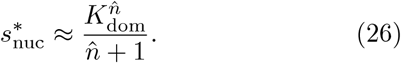

As in the equilibrium analysis in section II B, under optimal conditions nucleation proceeds nearly as if the total subunit density were amplified by the partition coefficient *K*_dom_.

### 3. Effect of LLPS on maximal assembly rates and kinetic traps

We now evaluate the effect of the domain on the propensity of the system to undergo the monomer-starvation kinetic trap, by evaluating the dependence of the elongation and median assembly timescales on the phase separation parameters.

The median assembly timescale can be computed by the same analysis used above for the nucleation time, leading to

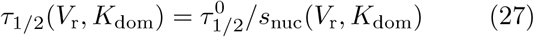

and similarly the threshold concentration below which nucleation does not occur on relevant timescales is

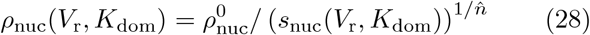

with 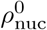 given by Eq. (22).

The elongation time within the domain, *τ*_elong,D_ or background *τ*_elong,B_ is given by Eq. (B1) with the appropriate local concentration 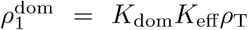 or 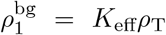. To estimate the onset of the kinetic trap, we must account for the numbers of capsids that are forming by both reaction channels (in the domain or background), so we compute an average elongation time weighted by the relative number of assemblies that form the domain or background

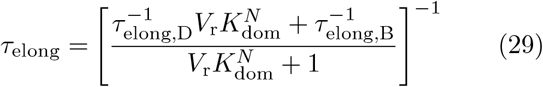

In the limit of strongly forward-biased elongation and 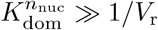 so all nucleation occurs in the domain, the elongation timescale is approximately

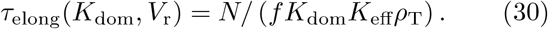

As discussed in section II D 1, the minimum assembly timescale occurs when the nucleation and elongation timescales are equal, *τ*_elong_(*K*_dom_, *V*_r_, *ρ*_*_) = *τ*_1*/*2_(*K*_dom_, *V*_r_, *ρ*_*_); the monomer-starvation kinetic trap begins occur beyond this point. Using Eqs. (25), (27), and (30) results in

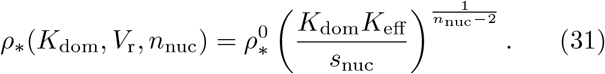

Finally, we can approximately extend the scaling estimates of this section (II D) to the CNT model by substituting Eq. (18) for *n*_nuc_.

Eq. (31) shows that a key feature of preferential partitioning into the domain is the ability of the system to buffer itself against the monomer-starvation kinetic trap while maintaining fast *localized* assembly in the domain, as shown in Fig. 4. We can further assess this feature in several ways as follows.

Fig. 6A,B compares Eqs. (27) and (29) to the median assembly and elongation times computed from the rate equations as a function of *ρ*_T_ and *K*_dom_ respectively. We see that the scaling estimates and numerical results closely agree until the nucleation and elongation timescales become comparable; the threshold concentration *ρ*_*_ (Eq. (31)) and partition coefficient 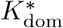 (Eq. (C3)) are shown as a vertical dashed lines in Figs. 6A,B respectively. Figs. 6C,D show the median assembly timescale assembly times computed from the rate equations as a function of *ρ*_T_ and *K*_dom_ and *V*_r_ respectively, with the locus of parameter values leading to minimum assembly predicted by Eq. (31) shown as white ◦ symbols. The prediction closely tracks the minimum assembly timescale observed in the numerical results. Below this threshold the median assembly time is closely predicted by Eq. (27), with the assembly timescale sped up according to *s*_nuc_ in Eq. (25). Above this threshold the numerically computed assembly timescales rapidly increase due to overly fast nucleation and thus onset of the monomer-starvation trap. We also show the CAC and the threshold for achieving assembly within an observation time of 1 day on these plots. Notice that, at a given value of *V*_r_, there is an optimal value of *K*_dom_ (estimated below) which maximizes robustness of assembly to variations in concentration. In contrast, robustness monotonically increases with decreasing *V*_r_.

**FIG. 6.**
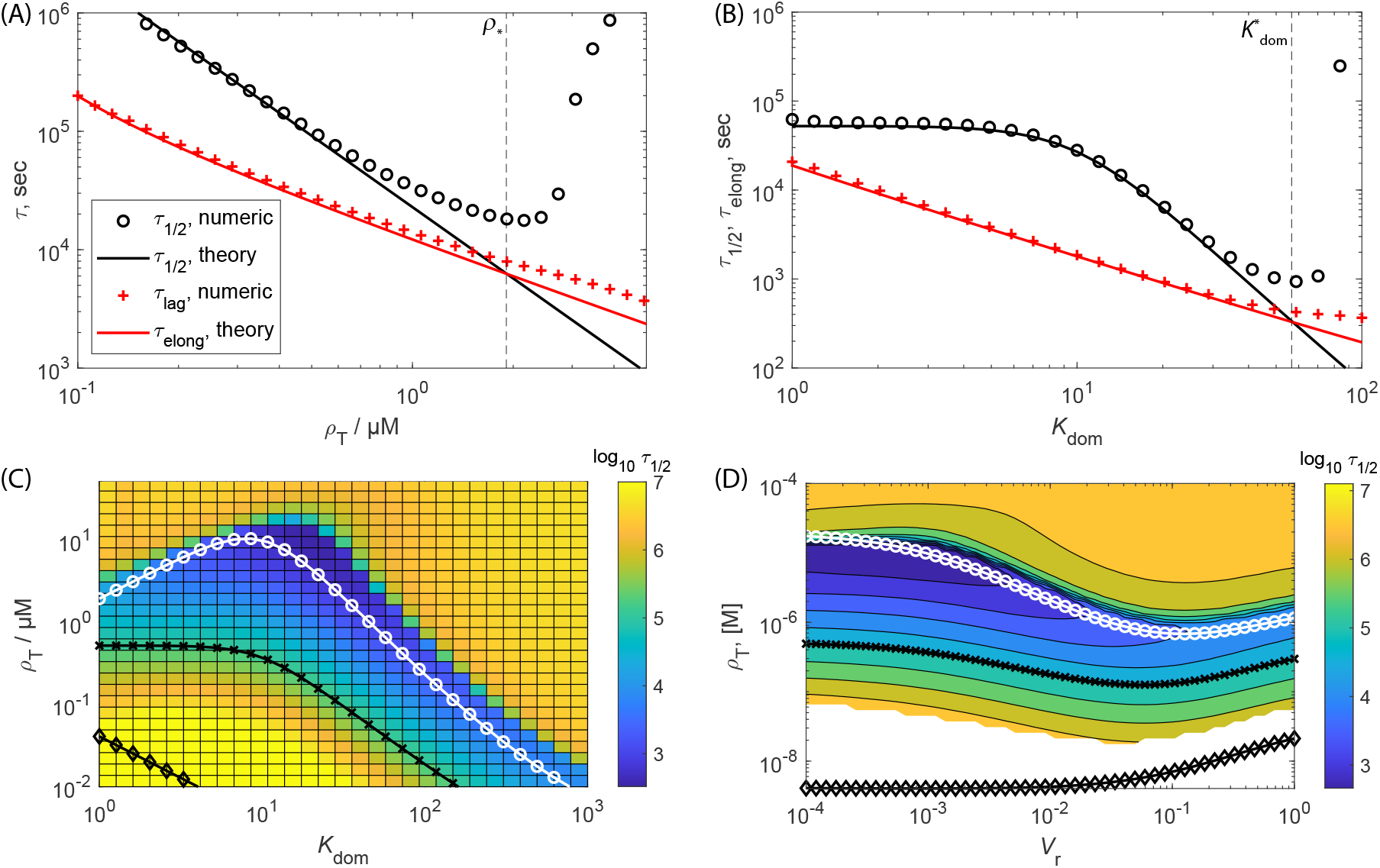
Effect of LLPS on assembly timescales and the monomer-starvation kinetic trap for the NG model. **(A)** The median assembly time *τ*_1*/*2_ and lag time calculated numerically from the Master equation (Eq. (13)) and scaling estimates for the median assembly time (Eq. (21)) and elongation time (Eq. (B1)) as a function of subunit concentration, with no LLPS. The vertical dashed line indicates the concentration corresponding to the onset of the monomer starvation kinetic trap (*ρ*_*_, Eq. (31)). **(B)** Same quantities, shown as a function of the partition coefficient for concentration *ρ*_T_ = 0.7*µ*M. The vertical dashed line shows the estimate of the optimal value for the partition coefficient, 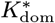 (Eq. (C3)). The domain ratio is *V*_r_ = 10^*−*3^ for (A) and (B). **(C, D)** The median assembly time predicted by the rate equation model as a function of subunit concentration and **(C)** varying domain partition coefficient with *V*_r_ = 10^*−*3^ or **(D)** varying *V*_r_ with *K*_dom_ = 10. The white line and ‘◦’ symbols correspond to the theoretical prediction for the relationship between the subunit concentration and partition coefficient corresponding to the minimal assembly timescale (Eq. (31)), beyond which the monomer-starvation kinetic trap begins to set in. The black line and ‘x’ symbols correspond to the relationship between the subunit concentration and *K*_dom_ value (Eq. (28)) below which nucleation will not be observed on an experimentally relevant observation timescale of *τ*_obs_ = 1 day. The black line and ‘♢’ symbols denote the concentration and *K*_dom_ values corresponding to the CAC (Eq. (9)). Other parameters are *N* = 120, *n*_nuc_ = 3, *g*_elong_ = −17*k*_B_*T*, and *g*_nuc_ = −4*k*_B_*T*.

#### The maximum assembly speedup depends on volume ratio, critical nucleus size, and subunit concentration

Given that the domain both shifts and broadens the range of parameter values over which productive assembly can occur, it is of interest to determine parameters for which LLPS has the strongest effect on assembly times. To this end, we define the assembly ‘speedup’ as the factor by which the median assembly time decreases with LLPS relative to bulk solution: *s*_LLPS_(*K*_dom_, *V*_r_) ≡ *τ* ^0^*/τ*(*K*_dom_, *V*_r_). Recalling that the minimum assembly timescale occurs at *ρ*_*_ when elongation and nucleation times are equal, we can then maximize the speedup with respect to the partition coefficient to obtain (appendix C)

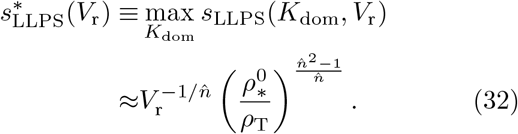

Thus, for an optimal domain partition coefficient, assembly can be sped up (i.e. *τ*_1*/*2_ reduced) by orders of magnitude for small *V*_r_. The degree of speedup increases with: decreasing *V*_r_, increasing critical nucleus size, and decreasing total subunit concentration. These trends can be understood as follows. Decreasing *V*_r_ means that subunits are not depleted as quickly within the domain, thus allowing larger values of *K*_dom_ and correspondingly higher local concentrations of subunits within the domain without depleting subunits quickly enough to cause over-nucleation and monomer starvation. A larger critical nucleus size provides a larger separation between nucleation and growth timescales, thus enabling further concentration of subunits in the domain without overnucleation. The decreasing dependence on concentration arises because as the system approaches 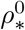, the assembly timescale without LLPS decreases and thus so does the extent of possible speedup before over-nucleation sets in. However, note in Fig. 7 that the maximum optimal concentration in the presence of LLPS exceeds the intrinsic value, 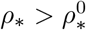, due to the extra regulation of nucleation and growth timescales allowed by a domain. Also note that LLPS provides speedup even after monomer starvation begins to set in.

**FIG. 7.**
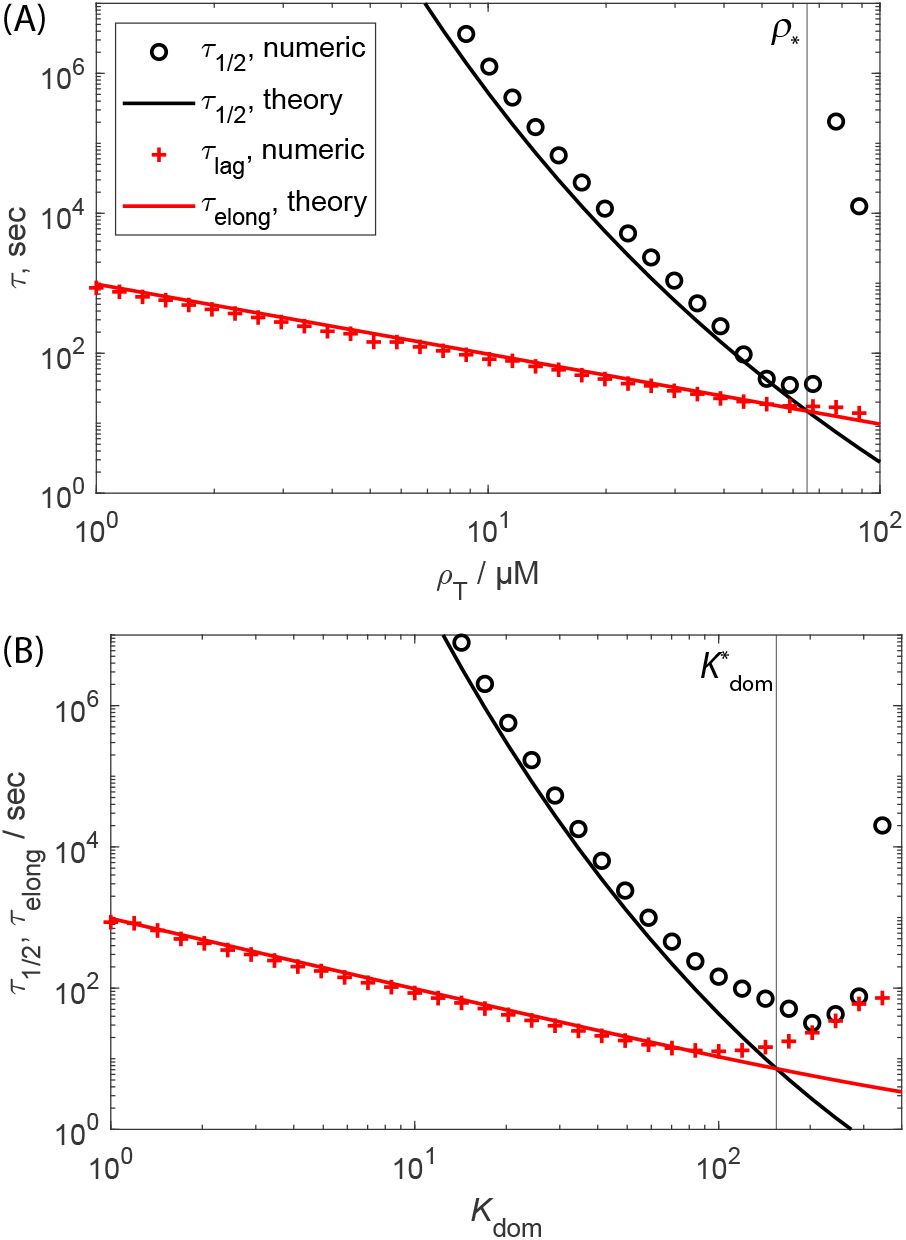
Effect of LLPS on assembly timescales and the monomer-starvation kinetic trap for the CNT model. **(A)** The median assembly time *τ*_1*/*2_ and lag time calculated numerically from the Master equation (Eq. (13)) and scaling estimates for the median assembly time (Eq. (21)) and elongation time (Eq. (B1)) as a function of subunit concentration, with no LLPS. The vertical dashed line indicates the concentration corresponding to the onset of the monomer starvation kinetic trap (*ρ*_*_, Eq. (31)). **(B)** Same quantities, shown as a function of the partition coefficient for concentration *ρ*_T_ = 0.6*µ*M. The vertical dashed line shows the estimate of the optimal value for the partition coefficient, 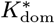 (Eq. (C3)). Parameter values are *N* = 120, *g*_sub_ = −17*k*_B_*T*, and *V*_r_ = 10^*−*3^.

Fig. 8 compares the scaling estimate for speedup (Eq. (C4)) to the value computed numerically from the rate equations. For the numerical value, we computed 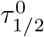 for fixed *ρ*_T_ and interaction parameters by numerically integrating the rate equations without LLPS, and then performed numerical minimization over *K*_dom_ to obtain 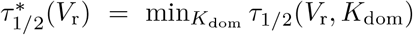 with respect to *K*_dom_ for the same *ρ*_T_ and interaction parameters. Then the speedup is given by 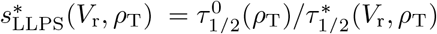. We have presented the speedup as a function of concentration normalized by the optimal value in the absence of LLPS so that the results can be shown on the same plot. As shown in Fig. 8A, the scaling estimate closely matches the numerical result until *ρ*_T_ exceeds the maximum value of *ρ*_*_ at which point monomer starvation begins to set in. The results for the CNT model are obtained by substituting Eq. (18) into Eqs. (C2))-(C4). The agreement is reasonable but not as close as the NG estimate because Eq. (18) is based on the critical nucleus size in the absence of LLPS. As noted above, we see that LLPS continues to speedup the assembly time even in the monomer-starvation regime.

**FIG. 8.**
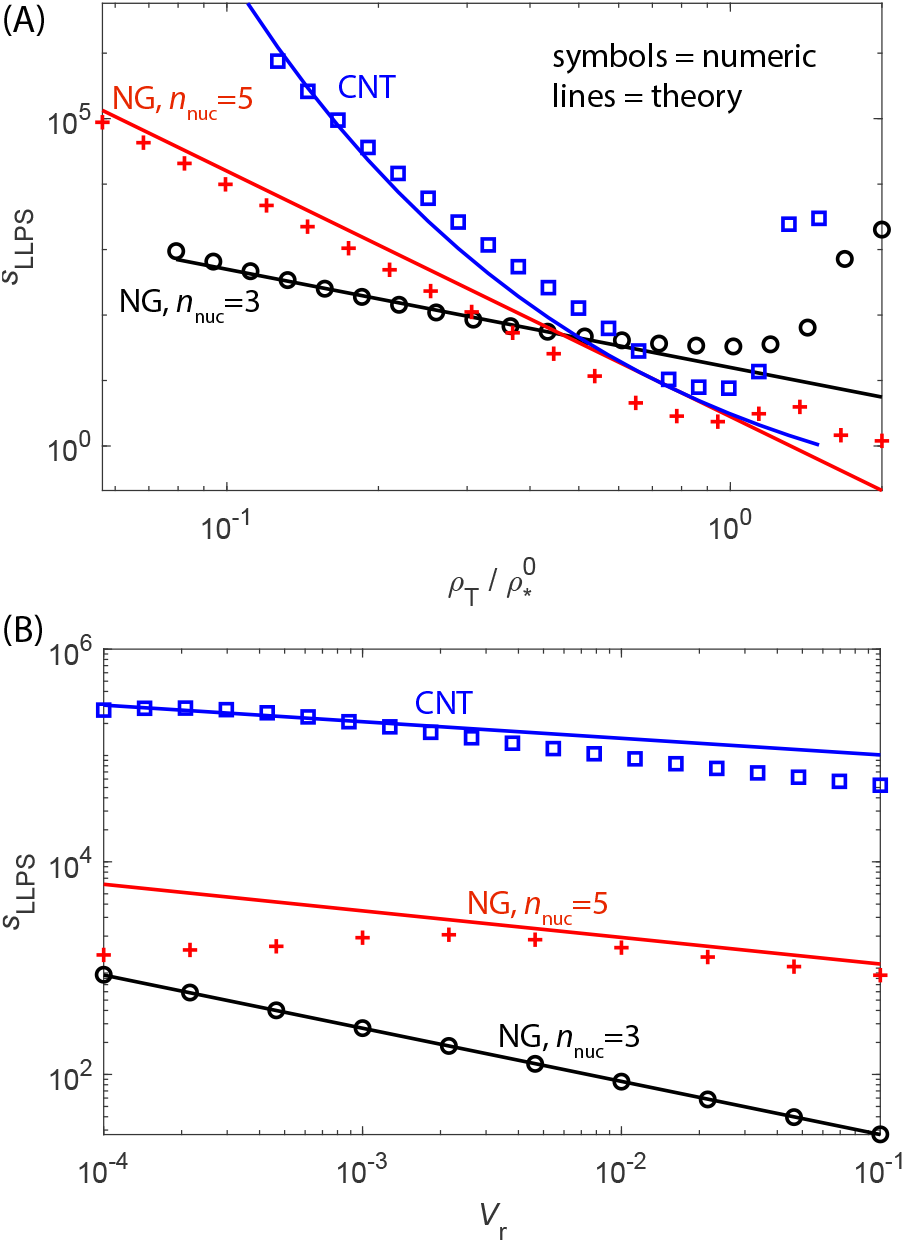
Maximum speedup provided by LLPS accounting for kinetic trapping. **(A)** The assembly speedup optimized over the domain partition coefficient, 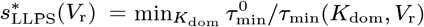 is shown as a function of subunit concentration *ρ*_T_ for fixed *V*_r_ = 10^*−*3^. Results are shown for the NG model with critical nucleus sizes *n*_nuc_ = 3 and *n*_nuc_ = 5, as well as the CNT model. The symbols show results obtained from the Master equation with τ _min_ calculated by numerically minimizing *τ*_1*/*2_ with respect to *K*_dom_. The lines show the approximate estimate Eq. (C4). The subunit concentrations are scaled by the optimal concentration in the absence of LLPS, 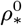, so that the results are visible on a single plot. The optimal concentrations for these parameters are 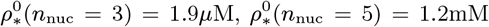 for the NG model and 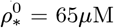 for the CNT model. **(B)** The assembly speedup optimized over *K*_dom_ as a function of *V*_r_ for fixed subunit concentration *ρ*_T_*/ρ*_*_ = 0.15.

Fig. 8B shows the speedup as a function of *V*_r_ for fixed *ρ*_T_. Here we see good agreement between the scaling estimate and numerical results, except the numerical speedup diminishes at small *V*_r_ for *n*_nuc_ = 5. This occurs because the minimum assembly timescale has decreased below the diffusion limited timescale (Eq. (A4)) in this regime, which is not accounted for in the scaling estimate.

#### Maximizing assembly robustness

Notice that 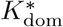 decreases with increasing subunit concentration (Eq. (C2)). For large subunit concentrations (i.e. 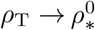, the assumption that assembly occurs primarily in the domain breaks down and we must consider the full form of *s*_nuc_ (Eq. (25)). If we substitute this into the expression for Eq. (31), we see that there is a maximum in *ρ*_*_ at

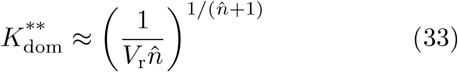

which results in

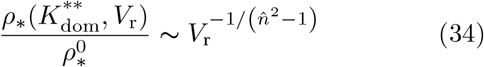

Finally, using Eq. 9 shows that the range of subunit concentrations over which assembly is favorable increases as

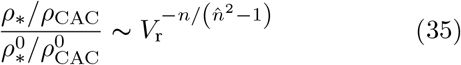

Importantly, for small *V*_r_ the width of this range, and thus the robustness of assembly to variations in subunit concentration or subunit-subunit binding affinities, increases by orders of magnitude (Fig. 6D).

We can alternatively specify robustness by defining the region of *productive assembly* as the set of parameter values for which nucleation occurs within experimentally relevant timescales (e.g. 1 day) and avoids the monomer-starvation trap. To maximize the breadth of this range, we define 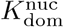 as the partition coefficient that maximizes the ratio of the monomer-starvation threshold to the nucleation timescale threshold (Eq. (28)): 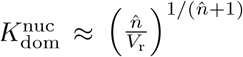

## III. CONCLUSIONS

It is well-established that efficient self-assembly in homogeneous solution is constrained to a narrow window of moderate subunit concentrations and interaction strengths, due to the competing constraints of minimizing nucleation timescales while avoiding kinetic traps [28–47]. Here, we find that when subunits preferentially partition into nanoor microscale domains, the range of parameters leading to productive assembly can be broadened by more than an order of magnitude, and the corresponding assembly timescales can be reduced by multiple orders of magnitude. Moreover, in part of this parameter range, almost all assembly occurs within the domain interior, thus allowing spatial control over assembly. These behaviors depend sensitively on two parameters that control phase coexistence: the partition coefficient of subunits into phase separated domains and the size ratio between the domains and the cell. In addition, we find that the maximum degree of speedup due to LLPS increases with: decreasing domain/cell size ratio or subunit concentration and increasing assembly critical nucleus size. These effects arise because the domain (or domains) drive high local concentrations of subunits, thus minimizing the local nucleation timescales, but the small size of the domain limits the total nucleation rate (averaged over the whole system volume). In effect, the bulk exterior acts as a subunit ‘buffer’ that, early in the reaction, steadily supplies subunits to the domain and thereby suppresses the monomer starvation kinetic trap (see Fig. 4). This mechanism has the strongest effect on robustness of assembly to variations in parameter values for small critical nucleus sizes or non-nucleated reactions, for which the homogeneous system lacks an intrinsic difference between nucleation and growth timescales and thus is most sensitive to subunit depletion. However, the decrease in assembly timescales is most dramatic for larger critical nucleus sizes, due to the high-order dependence of assembly timescales on local subunit concentration.

### Relevant parameter ranges

Since these mechanisms depend on localization of subunits, the ability of LLPS to control assembly increases with decreasing domain size (relative to the total system size). To estimate the relevance of this effect in biological systems, consider that typical domains in eukaryotic cells range in size from ∼50nm to 10 µm [54, 117, 118, 121]. For a domain with diameter 1 µm in a cell with diameter 20 µm, the volume ratio of the domain relative to bulk is *V*_r_ ∼ 10^−4^. From Eqs. (34)-(35) and Fig. 6, we see that the range of subunit concentrations leading to productive assembly could increase by up to two orders of magnitude, with increases in assembly rates exceeding five orders of magnitude (Eq. (C4) and Fig. 8). These increases reflect the ability of compartmentalization to enable fast localized assembly while minimizing the rate of global depletion of subunits. However, recall that the rate equation models and scaling estimates considered here do not account for kinetic traps resulting from malformed assemblies that can arise under overly rapid subunit association [11, 32, 39, 41, 138].

### Testing in experiments

Since our models are general, the quantitative predictions and scaling formulae described here could be tested in any experimental system for which there is phase coexistence, the assembly subunits preferentially partition into one phase, and the size of phase-separated domains can be controlled. While phase separation appears to be ubiquitous in biological cells, there is greater ability to control the size and composition of domains with *in vitro experiments* [52, 54, 76, 99]. Domain sizes can be controlled in bulk systems by varying the total density of the phase separating components, while microfluidic arrays enable precise control over droplet sizes and compositions.

### Outlook

We have focused on a minimal model for this first study of the effects of LLPS on assembly robustness. There are a number of additional physical ingredients that merit further exploration. For parameter regimes that lead to high subunit concentrations within the domain, the assumption that the subunits do not affect the equilibrium domain size and composition will break down. To be applicable at high concentrations the model should also allow for malformed assemblies [32, 39]. Further, Schmit and Michaels showed that the subunit diffusion slows with increasing subunit-domain attraction strength (*g*_dom_ in our model), then there is an optimum *g*_dom_ beyond which assembly slows. The results in our work arising from competing interactions are distinct from this effect. Other important effects to be incorporated include: slow diffusion into/out of the domain [129], accounting for spatial structure and stochasticity of assembly [32, 139–142], nonequilibrium effects such as synthesis of new subunits or phosphorylizationdriven changes in assembly activity, selective partitioning of different species within in a multicomponent assembly reaction, and the ability of the domain to template the size and shape of assemblies, such as occurs in bacterial microcompartments [43, 45, 123, 124]. Ultimately, understanding how different combinations of these physical mechanisms enable phase separation processes to control the time, place, and rate of assembly will engender a more complete understanding of biological self-assembly, and can advance strategies for designing human-engineered nanostructured materials.

## ACKNOWLEDGMENTS

This work was supported by Award Number R01GM108021 from the National Institute Of General Medical Sciences and the Brandeis Center for Bioinspired Soft Materials, an NSF MRSEC, DMR-2011846. Computational resources were provided by NSF XSEDE computing resources (Expanse) and the Brandeis HPCC which is partially supported by DMR-2011846. We gratefully acknowledge William Jacobs for insightful comments on the manuscript.

## Appendix A Kinetics of partitioning

In this appendix we consider the kinetics of subunits and assemblies partitioning between the domain and background. We will see that under most cases the rate of subunits partitioning into the domain will be fast relative to assembly timescales, and thus in subsequent sections we will develop scaling estimates based on a quasiequilibrium assumption for the ratio of subunit concentrations in the domain and background.

For a single spherical domain, the rate for intermediates of size *n* to enter the domain is given by 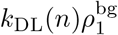, with the diffusion-limited adsorption rate constant *k*_DL_(*n*) = 4*πR*_dom_*D*_*n*_, with *R*_dom_ the radius of the domain, and *D*_*n*_ the diffusion constant for the intermediate. The dependence of diffusion constant on particle size is complex due to crowding and active processes (e.g. [143, 144]). The diffusion coefficient for individual proteins is typically ∼0.1*D*_0_, with *D*_0_ = 100*µ*m^2^*/s* the value in dilute solution. For simplicity, we assume a form

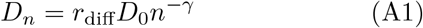

with *r*_diff_ the ratio of the diffusion coefficient for a protein monomer in the cell relative to dilute solution, and the factor *n*^*γ*^ gives the scaling of the hydrodynamic radius with size for the assembly, which depends on its geometry. In the results below we will typically set *r*_diff_ = 0.1 and *γ* = 1/2, where the latter is the scaling for a capsid-like assembly. In general the results will not depend (even quantitatively) on these values. However, if the diffusion of large intermediates and assembly products is sufficiently suppressed within a cell, the distribution of assembly sizes will be under kinetic control even at long times.

Finally, to allow for the possibility of arrested or microphase separation, we could consider *n*_dom_ domains with radius 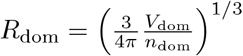

Putting these considerations together results in a diffusion limited rate constant

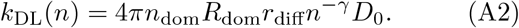

For simplicity we assume diffusion rates are uniform throughout the background and domain).

At equilibrium, the flux of subunit adsorption into the domain must be balanced by the outward flux, leading to the detailed balance condition on the desorption rate constant (*k*_desorb_) relative to adsorption, *k*_DL_*/k*_desorb_ = *K*_dom_. Thus, if we assume a fixed subunit concentration far from the domain 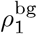, the kinetics of adsorption into the domain follows

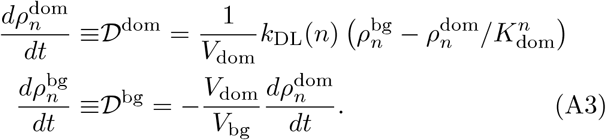

To get a feeling for the numbers, let us consider a single domain with radius *R*_dom_ = 1 *µ*m in a cell with radius *R*_cell_ = 10 *µ*m (so *V*_r_ = 10^−3^), and typical diffusion coefficient for protein in the cytoplasm *D* = 10 *µ*m^2^/s (*r*_diff_ = 0.1), which gives a diffusion-limited rate for monomers of *k*_DL_ = 4*π* × 10^10^ nm^3^/s = 7.6 × 10^10^/M·s. In the limit *f*_selec_ ≫ 1 so that essentially all assembly occurs within the domain, the characteristic timescale for a significant fraction of subunits to reach the domain is given from Eq. (14) as

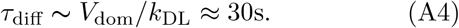

Under most conditions that we consider below this will be significantly faster than the characteristic assembly nucleation timescale and hence diffusion timescales can be neglected. However, under conditions of extremely strong partitioning into the domain and thus high local concentrations and rapid nucleation, the ability of a single domain to accelerate assembly will be limited by this diffusion timescale. But, allowing for multiple domains would reduce this timescale.

## Appendix B Elongation timescales

For the model we are focusing on, a linear nucleation and growth process with *f* independent of intermediate size, we can specifically estimate the elongation timescale by the mean first passage time for a biased random walk with a reflecting boundary conditions at *n*_nuc_ and absorbing boundary conditions at *N*, with forward and reverse hopping rates given by the subunit association and dissociation rates respectively [130]. For early in the reaction, when *ρ*_1_ ≈ *ρ*_T_, this results in

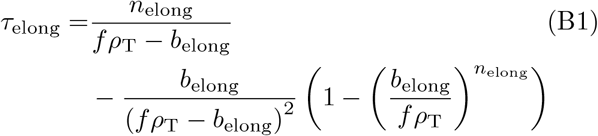

with *n*_elong_ = *N* − *n*_nuc_ ≈ *N*. In the limit of *fρ*_T_ ≫ *b*_elong_ Eq. B1 can be approximated to give *τ*_elong_ ≈ *n*_elong_*/fρ*_T_, or *α* = 1 in Eq. 19, while for similar forward and reverse reaction rates, *fρ*_T_ ≈ *b*_elong_, it approaches the solution for an unbiased random walk 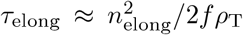, or *α* = 2. For the CNT model, since association rates are ∝ *n*^1*/*2^, we obtain *α* = 1/2 for strongly forward-biased assembly [130].

## Appendix C Maximum assembly speedup

In this appendix we evaluate the maximum possible speedup due to LLPS accounting for the monomer depletion kinetic trap. Recalling that the minimum assembly timescale *τ*_min_ occurs at *ρ*_*_ when the elongation and median assembly timescales are equal, we obtain

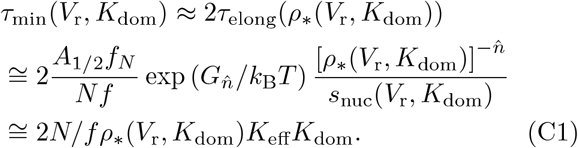

Defining the assembly speedup as *s*_LLPS_(*K*_dom_, *V*_r_) ≡ *τ*^0^ */τ*(*K*_dom_, *V*_r_), our goal will be to estimate the maximum speedup possible at a given subunit concentration and domain size by optimizing *τ*_min_ over *K*_dom_. To simplify the calculation, we assume that LLPS dominates near the maximal speedup and thus assembly primarily occurs in the domain, so we can simplify the nucleation acceleration as 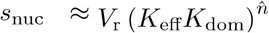, the elongation time as *τ*_elong_ ≈ 2*N/fρ*_*_*K*_eff_*K*_dom_, and the median assembly time as 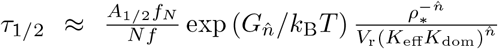 and 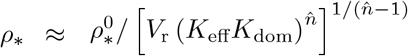.

Now, we optimize over *K*_dom_ to obtain

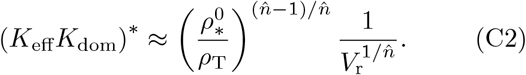

Solving for the optimal value of the partition coefficient 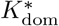 then yields

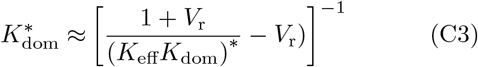

with 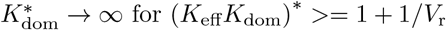 for (*K*_eff_*K*_dom_)^*^ >= 1 + 1*/V*_r_.

We then use Eq. C2 obtain the *maximum* speedup as

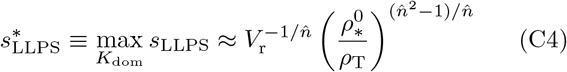

## Appendix D Analysis for *n*_nuc_ = 2

### Assembly timescales without LLPS

To this point we have assumed the existence of a nucleation barrier and thus a critical nucleus size *n*_nuc_ ≥ 3 in our expression for the dependence of the nucleation rate on subunit concentration Eq. (20). For the special case of no nucleation barrier (*n*_nuc_ = 2), the reaction is unregulated and reduces to the well-studied case of unregulated 1-D filament assembly. If the association rate is independent of assembly size *n*, assembly rapidly results in an exponential distribution of filament sizes whose mean gradually increases through coarsening (but is cut off by the maximal size *N*). Regulation can be restored if the association rate of the first two subunits is slow compared to subsequent association events (e.g. if the subunits must undergo a conformational change in order to assemble [137]). In this case, the median assembly time is given by 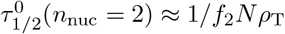 with *f*_2_ the dimerization rate constant. Equating 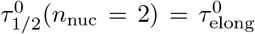 then shows that the form of the kinetics is independent of *ρ*_T_, but the reaction behaves as if regulated when *f*_2_ ≪ *f* with the threshold dimerization rate given by 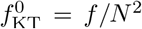. Here we have assumed that the association rate constant *f* is independent of size for *n >* 2 for simplicity.

### Assembly timescales with LLPS

Accounting for the presence of a domain results in

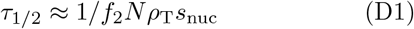

and *τ*_elong_ given by Eq. 30. As in the case without LLPS, the threshold for monomer-starvation is independent of total subunit concentration, but the threshold dimerization rate constant is now given by

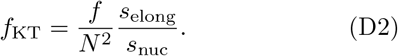

Importantly, Eq. (D2) shows that LLPS allows for the reaction to be regulated even for the case of the rate constant independent of assembly size (*f*_2_ = *f*).

